# Network-based analysis reveals microRNA regulation of oncogenic pathways in SOX10-depleted uveal melanoma

**DOI:** 10.64898/2025.12.05.688852

**Authors:** Chunyan Luan, Anja Wessely, Zhesi Zhang, Liang Zhang, Adrian Weich, Christopher Lischer, Carola Berking, Markus V. Heppt, Julio Vera, Xin Lai

## Abstract

SOX10 is essential for melanocyte development and maintenance and plays a critical role in uveal melanoma (UM) initiation and progression. While SOX10’s transcriptional regulation of protein-coding genes is well characterized, its role on microRNA (miRNA) regulatory landscape in UM remains unexplored. Here, we employed network-based modeling to systematically characterize miRNA regulatory functions following SOX10 depletion in UM. First, we profiled mRNA and miRNA expression levels in SOX10 wild-type and knockdown UM cells. Then, we integrated the transcriptomic data, a UM network, and a Bayesian model to quantify miRNAs’ regulatory activities and identify key miRNAs. Subsequently, we employed pathway enrichment analysis combined with literature mining to elucidate the functional roles of identified miRNAs through their target genes and associated signaling pathways in UM. We identified 17 miRNAs that show significant changes in regulatory activities following SOX10 knockdown in UM cells. These miRNAs regulate the expression of genes involved in cancer hallmark pathways, including cell cycle progression, mTORC1 signaling, and fatty acid metabolism. Notably, miR-34a, miR-25, miR-186, and miR-211 have tumor-suppressive potential by targeting genes involved in UM progression and metastasis. Our results suggested that SOX10 depletion in UM can activate tumor-suppressive mechanisms through regulating miRNAs.

## INTRODUCTION

Uveal melanoma (UM), the most common intraocular tumor in adults, arises from melanocytes in the choroid, ciliary body, or iris (1), but is far rarer than cutaneous melanoma (CM) and classified as an orphan cancer (2). Despite both tumors originating from melanocytes, UM and CM differ in tumor biology, clinical characteristics, and treatment options (3). An example of these differences is SOX10, a member of the SOX family of transcription factors that plays a critical role in neural crest development and the differentiation of various neural crest derivatives, including the melanocyte lineage (4–6). While SOX10 has been extensively studied in CM and is recognized for its crucial role in promoting tumor progression in melanoma (7–9), the knowledge about its function in UM remains limited, with a few studies showing that SOX10 has similar functions across UM lineages, such as strong nuclear expression regardless of their origins (10). SOX10 is expressed diffusely across 11 UM cell lines and in surrounding uveal or choroidal melanocytes (11). While SOX10 expression is typically consistent, a pathogenic SOX10 mutation is identified in a single UM patient, suggesting the existence of rare genetic subtypes with distinct biological characteristics (12). Mechanistically, SOX10 ablation, downregulating the expression of GAPDH, significantly suppresses UM growth and proliferation *in vivo* (13). BAF inhibition leads to the loss of enhancer occupancy by SOX10 and its interacting partner MITF, resulting in down-regulation of the melanocytic gene expression program in UM (14). In clinical applications, SOX10 serves as a critical diagnostic marker, exhibiting exclusive nuclear positivity in all 38 cases of a UM cohort and representing a sensitive marker for UM diagnosis (10).

SOX10 is a critical regulator in tumor progression, with its multifaceted functions involved in cell apoptosis, proliferation, and migration (15). Its regulatory landscape is further intricated by other molecules, such as microRNAs (miRNAs), which are a class of non-coding RNAs with 22-25 nucleotides in length. By modulating the transcription and translation of specific protein-coding and non-coding genes such as circular RNAs, miRNAs can alter cellular behavior and influence the outcomes of various cellular processes alone or in a combined manner (16–21). Growing evidence suggests that miRNAs play a crucial role in the development and progression of UM (22–24). They act as vital regulators of complex signaling networks by targeting key genes involved in tumor growth, invasion, and metastasis (23, 25, 26). For example, the miR-181 family acts as a negative regulator of CTDSPL-mediated cell cycle progression (27). The long non-coding RNA ZNF197-AS1 also inhibits uveal melanoma cell proliferation, migration, and invasion by sponging miR-425, which upregulates GABARAPL1 (28). Additionally, miR-34a suppresses the migration and invasion of these cells through its negative control of LGR4 (29). Furthermore, miRNAs hold significant potential as diagnostic biomarkers (30–33) and therapeutic agents, either as monotherapy or in combination treatments, to target and modulate gene expression in cancer such as UM (34–36). miRNA-mediated regulatory circuits are crucial for tumor plasticity by controlling, for example the dynamics of SOX10 expression levels (37). Though SOX10-deficient CM cells have been reported to be associated with therapeutic resistance (38) and invasive phenotypes (39), there has been no investigation into altered miRNA profiles in UM, which prevents our understanding of the SOX10-mediated miRNA landscape in this malignancy. To address this knowledge gap, we investigated SOX10-mediated miRNA regulation and their regulatory profiles in UM. We hypothesized that SOX10 modulates specific miRNAs to coordinate downstream pathways involved in UM progression. Identifying key miRNAs that regulate tumor growth and proliferation may therefore provide deeper insights into SOX10’s mechanistic role in UM pathogenesis.

Here, we present a network-based approach to identify miRNAs regulated by SOX10 in UM (Figure 1). With this approach, we aim to characterize SOX10-mediated miRNA expression and regulatory profiles for better an understanding of the molecular function of miRNAs in UM. Specifically, we performed high-throughput RNA sequencing to obtain genome-wide transcriptomic profiles (including protein-coding genes and miRNAs) in SOX10-knockdown (SOX10-KD) UM cells. Next, we integrated the gene expression data with a UM network to quantify the regulatory activity of miRNAs. Then, we identified driver miRNAs that show significant changes in the regulatory activity of SOX10-KD UM cells, and we elaborated the molecular mechanisms through which these miRNAs can influence tumor phenotypes at the pathway level. Taken together, we demonstrated the power of network medicine approaches to identify functional miRNA-gene interactions with therapeutic potential in UM.

**Figure 1:**
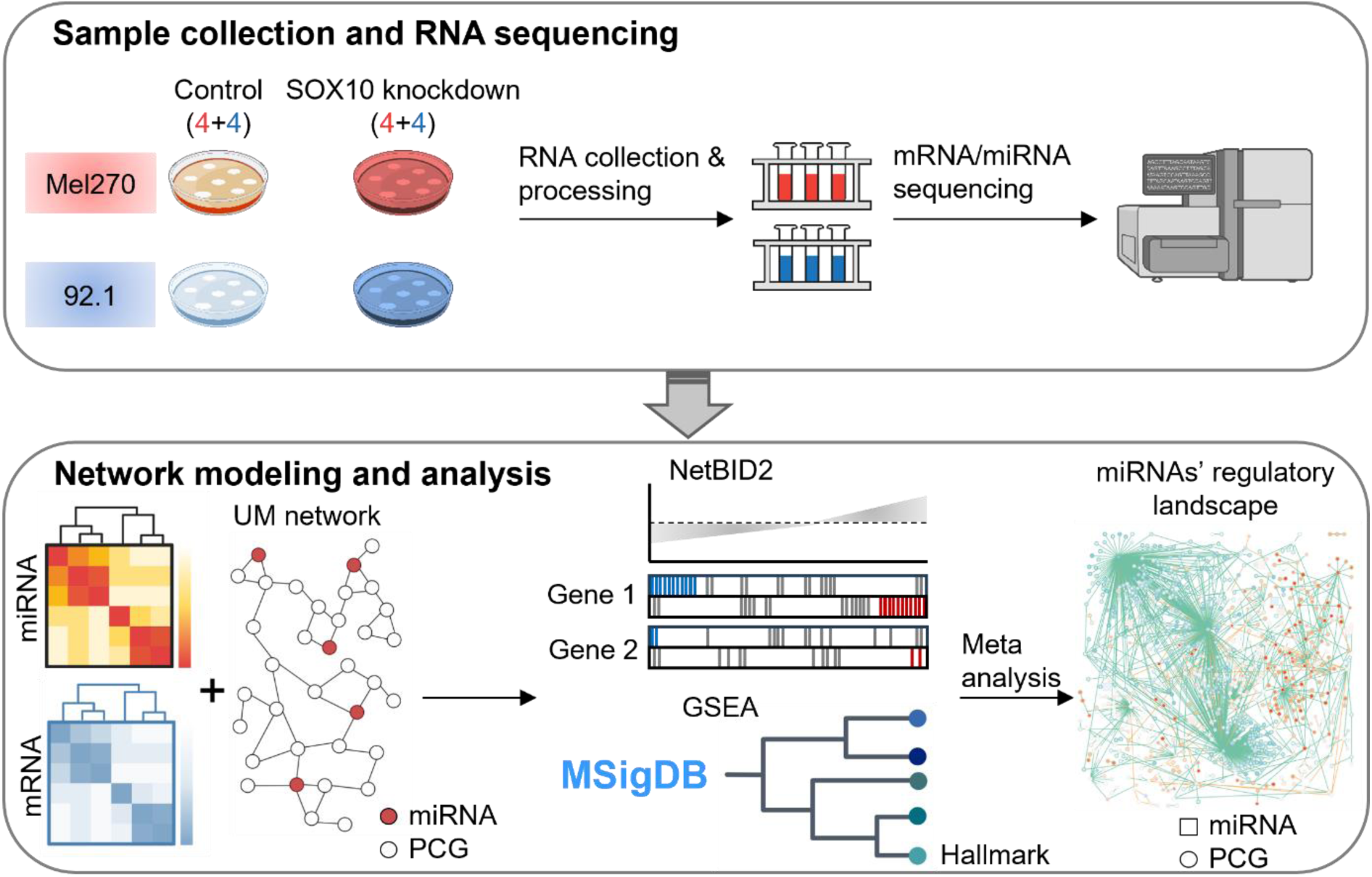
A network-based approach to investigate miRNA function in SOX10-KD UM cells. We began with the preparation of UM cells, where SOX10 is knocked down in the case group while the control group remains untreated. Following sample collection, we performed RNA sequencing to profile miRNA and mRNA expression levels. Subsequently, we input the data together with the network into NetBID2 that combines gene expression levels and miRNA-gene interactions to identify driver miRNAs. Finally, we performed gene set enrichment analysis (GESA) using the identified driver miRNAs’ targets to elaborate their roles in UM with the help of the UM network and literature mining. The detailed computational workflow is shown in Figure S1.

## MATERIALS AND METHODS

### Experimental design and cell culture

We used two UM cell lines (92.1 and Mel270) to investigate gene expression profiles (including mRNA and miRNA) under two experimental conditions: SOX10 knockdown (SOX10-KD) and control. The experimental design included four biological replicates for each cell line under each condition, resulting in a total of 16 samples.

UM cell lines 92.1 (RRID:CVCL_8607) and Mel270 (RRID:CVCL_C302) were kindly provided by Klaus Griewank, University Hospital Essen, Germany. The cells were cultured in RPMI1640 medium containing 2 mM L-glutamine (Gibco by Life Technologies, Waltham, Massachusetts, USA), 10% fetal bovine serum (FBS; Merck, Darmstadt, Germany), and 1x antibiotic-antimycotic (Invitrogen, Waltham, Massachusetts, USA) in a humidified incubator at 37°C and 5% CO2 atmosphere. The cells were regularly tested for mycoplasma contamination using the Venor®GeM Classic Mycoplasma Detection Kit for conventional PCR (Minerva Biolabs, Berlin, Germany) according to the manufacturer’s instructions.

### siRNA transfection

For SOX10-KD, cells were seeded in 6-well plates in RPMI1640 medium containing 2 mM L-glutamine and 10% FBS without 1× antibiotic-antimycotic supplement and transfected the following day with 20 nM siRNAs: siSOX10-B (sequence 5’-GUAUGCAGCACAAGAAAGATT-3’) (40) or control siRNA with a random sequence that does not match the human genome (sequence 5’-GCGCAUUCCAGCUUACGUA-3’ with asymmetric dTdT overhang at the 3’ end; both from Eurofins Genomics, Ebersberg, Germany) using 1.25 µl Lipofectamine RNAiMAX (Invitrogen) per well. All cells were cultivated at 37°C and 5% CO2 atmosphere in a humidified incubator for 24 h.

### Total RNA extraction and qPCR to test efficacy of SOX10 knockdown

Total RNA was extracted 24 h after transfection using the miRNeasy Tissue Cells Advanced Kit including a DNase digestion step to eliminate genomic DNA contamination using the RNase-Free DNase Set (Qiagen, Hilden, Germany). RNA concentration was determined with a Nanodrop 2000 spectrophotometer (Thermo). SOX10-KD efficacy was assessed by quantitative real-time PCR (qPCR) before sequencing. Therefore, 1000 ng of total RNA were transcribed to cDNA using the Expand™ Reverse Transcriptase Kit (Roche, Penzberg, Germany) and oligo(dT)-oligonucleotide primers (Eurofins Genomics, Ebersberg, Germany). The LightCycler® TaqMan® Master Kit (Roche), 200 nM of forward and reverse primers designed with the software “Assay Design Center” (Roche) (Table S1), 100 nM primer-matched hydrolysis probes from the Universal ProbeLibrary set human (Roche), and 50 ng cDNA were used to carry out the qPCR on a qTower3 instrument (Analytik Jena, Jena, Germany). Gene expression was normalized to glyceraldehyde 3-phosphate dehydrogenase expression (GAPDH).

### RNA sequencing data processing and analysis

RNA sequencing analysis of pooled total RNA derived from four individual experiments was performed by CeGaT (Tübingen, Germany) using the KAPA RNA HyperPrep Kit with RiboErase (Roche) for library preparation and a NovaSeq™ 6000 instrument (Illumina, San Diego, California, USA) for sequencing. RNA quality was checked on a Bioanalyzer 2100 (Agilent Technologies, Santa Clara, California, USA) before RNA sequencing.

#### Quality control and quantification of mRNA and miRNA sequencing data

In our previous work, we used a standard bioinformatics pipeline to process mRNA sequencing data (41). Specifically, the raw sequencing data in FastQ format underwent quality control with FastQC (version 0.11.9) and trimming to remove adapters using TrimGalore (version 0.6.6). Then, the obtained reads were mapped to the human reference genome assembly GRCh38 (hg38) with Gencode Annotation (version 22) (42) by STAR (version 2.0201) (43). Finally, transcript quantification was made using StringTie (version 2.1.4), which accurately estimates the read counts for each transcript.

We used MultiQC to check the quality of the miRNA sequencing data (44) (Table S2). The sequencing data underwent comprehensive quality checks, including evaluation of base quality via Phred scores, assessment of sequence duplication levels to detect potential contamination or amplification bias, and analysis of sequence integrity after adapter removal. Additional quality control measures included examination of GC content and distribution of unspecific bases (or N bases) to identify possible sequencing biases. Then, we trimmed the reads by removing adapters using Trimmomatic (version 0.39) and retained good reads using fastq_quality filter with the parameters (-Q33 -q 20 -p 80). The trimmed reads were mapped to the human genome hg38 using STAR (version 2.7.3a) using the default parameters. After the alignment, we used samtools to convert and sort the sequence alignment/map files. Finally, miRNA read counts were quantified using FeatureCounts (version 2.0.1) using the human genome hg38.

#### Gene expression analysis

For gene expression analysis, Ensembl IDs were converted to gene symbols using the GRCh38 and miRNAs were named using mature miRNA names from miRbase (45). After the conversion, duplicate gene symbols were resolved by adding their expression values. We eliminated the batch effects of both mRNA and miRNA samples using *ComBat* in the R package *sva* (46). It is an empirical Bayesian method that can effectively adjust batch effects while retaining biological variation in the data. We performed principal component analysis (PCA) to visualize the results (Figure S2).

### Network modeling and analysis using NetBID2

We used NetBID2 to perform network modelling and analysis. It is a method that uses a network-based Bayesian inference method to identify drivers and shows its power to identify crucial genes with mild changes in their expression profiles (47, 48). The method is to identify drivers in gene regulatory networks, such as transcription factors (TF), signaling proteins, and miRNAs. In our case, the identified miRNAs are regarded as important regulators in a SOX10-mediated UM network. We merged the miRNA and mRNA expression profiles (Table S3) and obtained an overview of the data (Figure S3). Next, we normalized the data and performed a log2 transformation. The normalization method adds one to each gene’s raw count and multiplies it by a scaling factor, which is the mean sequence depth of all samples divided by the sum of the sequence depth of the corresponding sample. Finally, we filtered out genes whose expression values are at or below the fifth percentile in 90% or more of the samples, indicating that they are essentially not expressed in most samples.

#### The UM network reconstruction

We reconstructed a SOX10-centered gene regulatory network using annotated information containing the most relevant intracellular signaling pathways in UM through literature survey. The network was developed based on SOX10 interacting protein and further expanded and pruned using databases and available gene expression data for UM (see SM for details)(41). As a result, we obtained a SOX10-centered UM network with 2, 149 nodes (including 28 miRNAs) and 7, 832 edges (SM Excel S1). As the 16 UM samples’ mRNA and miRNA profiles are not paired, we divided them into four groups using matching cell lines (i.e. 92.1 and MEL270) and experimental conditions (control and SOX10-KD) (Table S3). We used the average expression levels of the four groups to compute Spearman correlation coefficients for interacting genes. Since the primary function of miRNAs is to repress gene expression in mammals (49), in our network we considered miRNA-mRNA interactions with negative correlation values as relevant and removed the interactions with positive values. The resulting regulatory network data, together with gene expression profiles, were used as inputs for performing differential gene expression (DE) and differential activity (DA) analysis.

#### Differential Expression and Activity inference

We used NetBID2, which uses a Bayesian linear modelling approach, for performing DA and DE analysis between SOX10-KD and control samples (48). Differentially expressed genes with an adjusted p-value ≤ 0.05 and |log2FC| ≥ 1 are considered significant. In our analysis, a gene’s activity is high when its up or downregulation manifests in the over or underexpression of its target genes. We used this method to infer miRNAs’ activity based on the correlation between the expression profiles of miRNAs and their targets. This analysis allowed us to quantify the functional impact of a miRNA within the regulatory network by measuring its downstream repressive effects rather than relying solely on its own expression level. The resulting activity score can be considered a metric of the miRNA’s regulatory influence within SOX10-KD UM cells. Mathematically, the activity score of a driver gene is computed using the following equation:

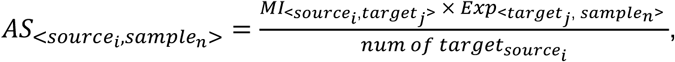

Where *MI_<i, j>_* is a matrix representing mutual information between source gene *i* (such as miRNAs), and target gene *j* (such as miRNA targets). *Exp_<j, n>_* is a matrix representing the expression of target *j* in sample *n*. The multiplication of the two matrices results in a score for source gene *i* in sample *n*, which is calculated by summing the expression levels of all targets in sample *n*, weighted by their mutual information with the specific source *i.* The source gene’s activity score is normalized by the number of the source gene’s targets. The sample-based genes’ activity scores (*AS_<i, n>_*) are used to conduct DA analysis between SOX10-KD and control samples using a Bayesian linear model. Genes with an adjusted p-value ≤ 0.05 are considered to have significant change in DA. The full results for DE and DA analysis can be found in SM Excel S2 and S3, respectively.

### Gene set enrichment analysis

We used Enrichr (50) to conduct gene set enrichment analysis for the 50 hallmark gene sets from MSigDB (51). The hallmark collection represents a refined, curated subset of gene sets designed to summarize well-defined biological states or processes with coherent expression patterns. For the background gene list, we used all genes in the UM network. For the input gene list, we used target genes of miRNAs with significant change in DA (adj. p-value ≤ 0.05). Gene sets with an adjusted p-value ≤ 0.05 were considered significant. In addition, the results included the pathways’ odds ratio and combined score (i.e., −log10(adj. p-value) multiplied by Z-score), where Z-score is a measure of the “deviation from the expected range” or the “deviation from the expected number of overlapping genes”. This combined score is related to the degree of enrichment, considering the size of the gene set and the total number of genes. The full results can be found in SM Excel S4.

## Results

### Integration of miRNA and mRNA data

We performed comprehensive quality control to check the reliability of miRNA sequencing data measured in UM cell lines. Although the data show good quality reflected by Phred Scores, we observed some abnormal and unusual incidences known to be common for miRNA sequencing data (Table S2). Specifically, the data has a high GC content, which skews the overall GC distribution. Due to the short size of miRNAs, the data shows a narrow sequence length distribution. In addition, certain miRNAs are usually highly abundant in cells (e.g., let-7 family miRNAs), leading to a higher proportion of miRNA overrepresented sequences compared to mRNAs (52). The overall diversity of miRNAs is generally lower than that of total RNA, leading to an increased proportion of duplicated sequences. However, the comprehensive quality check suggests that our miRNA data is of good quality and in line with data from other melanoma cell lines (53, 54).

We also generated comprehensive RNA sequencing data from matched SOX10-KD experiments in the same UM cell lines (i.e. 92.1 and Mel270) (41). Thus, we merged these complementary miRNA and transcriptomic datasets to investigate the regulatory network affected by SOX10-KD in UM cells, with particular emphasis on identifying miRNA-mediated mechanisms of gene regulation (Figure 1).

Following the data integration, we conducted quality control analyses to identify potential confounding factors. We used principal component analysis (PCA) to assess data heterogeneity and detected pronounced sample clustering based on cell line origin in the integrated gene expression profiles, indicating substantial batch effects that could obscure biological signals of interest (Figure S2). To preserve the SOX10-KD effect while minimizing technical variation, we applied a linear mixed-effects model to remove cell line-associated batch effects while retaining biologically relevant variation. Subsequent PCA validation confirmed the elimination of cell line-dependent clustering patterns while maintaining experimental group separation based on SOX10 status (Figure S3A), ensuring that subsequent comparison analysis reflects actual biological differences.

Furthermore, we found that the combined mRNA and miRNA expression profiles can distinguish SOX10-KD from control samples, with 6 out of 8 SOX10-KD samples clustering together and the remaining two clustering with control samples based on their expression profile distances (Figure S3B). All samples had many genes expressed at low levels (Figure S3C) and exhibited high correlation (Figure S3D). SOX10-KD resulted in massive change of gene expression, consistent with SOX10’s role as an important TF that regulates numerous genes in the melanocytic lineage and whose deregulation can substantially alter the transcriptomic landscape, as demonstrated in melanoma (39).

### SOX10 knockdown leads to massive transcriptomic change in UM

To characterize the effects of SOX10-KD in UM cells, we performed differential gene (DE) expression analysis. We identified 2, 222 up-regulated and 1, 156 down-regulated protein-coding genes with at least 2-fold changes (Figure 2A and SM Excel S2). The most significantly up-regulated genes (based on adjusted p-value) are BMF, FOXO1, and HSPB7, while the most down-regulated genes are KPNB1, GNPDA1, and RIOX2. The upregulated genes are reported to be related to tumorigenesis and development of UM. Specifically, BMF acts as a key mediator of apoptosis in UM, and it is targeted by miRNAs like miR-221 and miR-125b in cancer (55, 56). Its downregulation, particularly in response to HGF/cMET signaling, is a major mechanism by which UM cells acquire resistance to MEK inhibitors (57). An *in vitro* study indicated that HGF promotes resistance to MEK inhibitors by increasing the protein expressions of BCL2L11 and BMF in UM (58). Somatic mutations in FOXO1 are associated with increased metastatic potential and poorer prognosis in UM patients (59), and the gene is known to regulate multiple miRNAs in the context of cancer (60). HSPB7 is found to be expressed in UM subclusters in single-cell RNA-seq experiments (61) but there is no direct experimental evidence detailing its mechanistic role in UM. In contrast, the downregulated genes involvement in UM pathogenesis and progression remains to be investigated, but they are known to play a role in other cancers, for example KPNB1 in melanoma (62), GNPDA1 in several cancers (63) and RIOX2 in lung and prostate cancers (64, 65).

**Figure 2:**
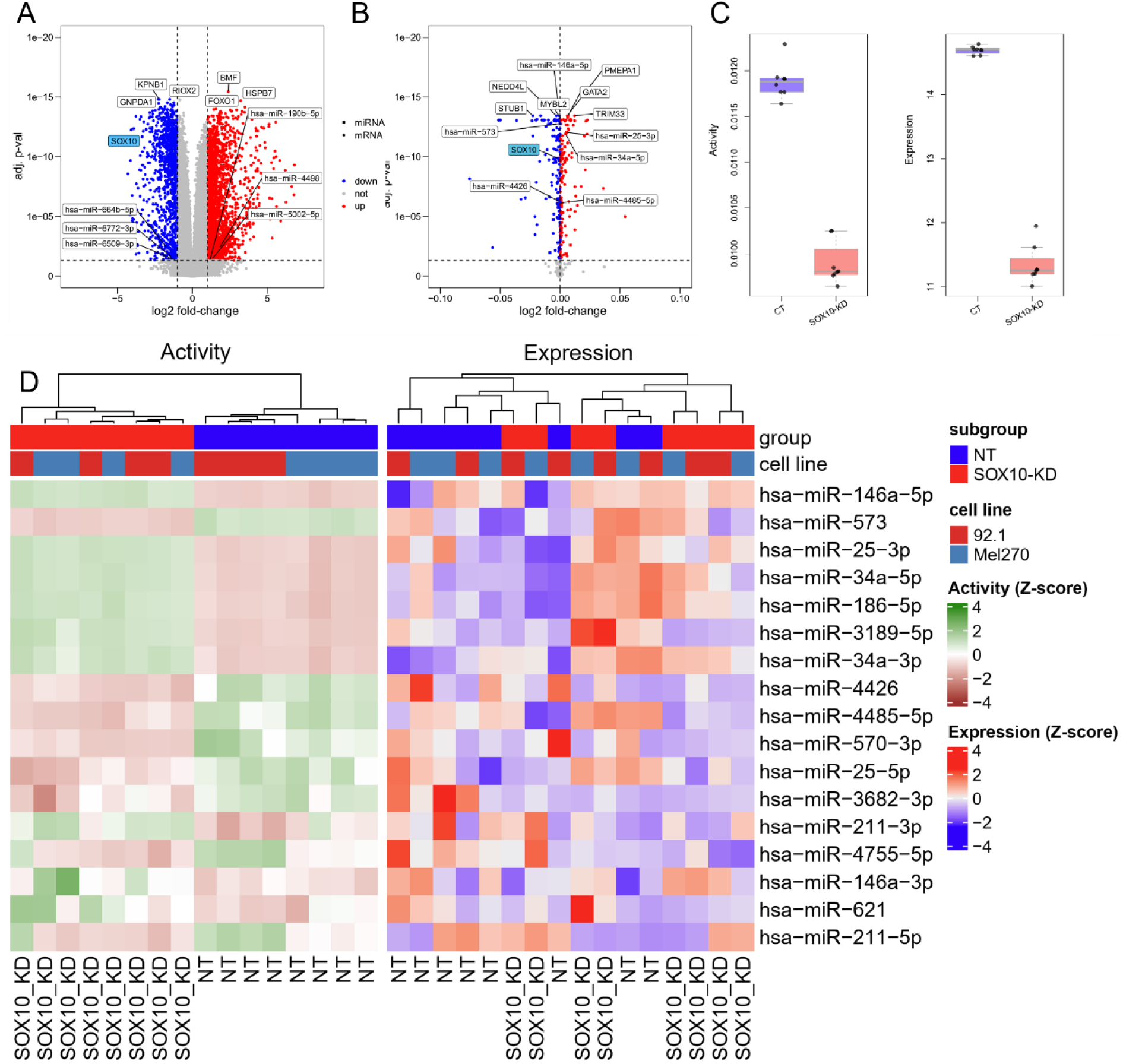
Differential expression and activity analysis. The volcano plots show differentially expressed genes (**A**) and genes with differentially regulatory activities (**B**) that are identified in SOX10-KD UM cells. Genes with significant change in gene expression (adjusted p-value ≤ 0.05 and |log2FC| ≥ 1) or regulatory activity (adjusted p-value ≤ 0.05) are shown in red (upregulated) and blue (downregulated), respectively. The labelled genes including protein-coding genes (square) and miRNAs (circle) are the top dysregulated ones (determined by adjusted p-value). SOX10 is highlighted in blue. (**C**) The box plot shows SOX10’s expression levels and regulatory activity in SOX10-KD and control groups (CT). (**D**) The heat map shows the expression levels and regulatory activity of the identified driver miRNAs. Both data are transformed using Z-scores. The samples are hierarchically clustered using the Euclidian distance between samples.

In addition, we identified six DE miRNAs including miR-664b-5p, miR-4498, miR-6772-3p that are significantly upregulated while miR-5002-5p, miR-6509-3p, and miR-190b-5p are significantly downregulated (Figure 2A). Though some of these miRNAs are involved in tumor progression, such as loss of miR-664b enhances malignancy of cutaneous melanoma (66) and miR-190b is a tumor suppressor in lung cancer (67), there is no evidence describing their association with UM.

Taken together, SOX10-KD in UM can massively alter transcriptomic landscape including mRNAs and miRNAs, suggesting its crucial role in regulating gene expression in UM. The results are consistent with previous findings that SOX10 is an important TF involved in cancer (15, 37) and other human pathologies (4).

### SOX10 knockdown leads to altered regulatory landscape in UM

Gene regulation encompasses a complex network of mechanisms that extend far beyond simple up- or down-regulation of mRNA or protein abundance (68, 69), so the change in expression levels does not fully reflect the importance of genes’ regulatory activities. Here, we used a network method to identify crucial genes, especially miRNAs, in SOX10-KD UM cells. Specifically, we first computed genes’ activity based on their expression profiles and regulatory interactions in a UM network developed by us and then we inferred genes with differential activities in SOX10-KD UM. The results show 168 and 174 protein-coding and miRNA genes with significantly upregulated and downregulated activities in SOX10-KD UM cells, respectively (Figure 2B and SM Excel S3). Among these genes, SOX10’s expression level and regulatory activity are significantly reduced in SOX10-KD UM cells consistent with our expectation (Figure 2C), demonstrating the method’s power to identify crucial genes in the UM network. As a transcriptional activator of many genes including TFs, SOX10-KD significantly downregulates several TFs’ expression in UM cells, including MITF (log2FC = −2.60; adjusted p-value = 2.95E-16), EDNRB (log2FC = −1.54; adjusted p-value = 1.81E-11), GJB1 (a.k.a. CX32; log2FC = −3.48; adjusted p-value = 2.1E-7), and MIA (log2FC = −1.21; adjusted p-value = 0.016). These alternations could drive a phenotype switch in UM. For instance, the disrupted SOX10-MITF pathway may lead to reduced cell proliferation, cell cycle arrest, and increased apoptosis (70, 71). The diminishment of transcriptional activity for key regulatory genes EDNRB (72, 73) and MIA (74) can affect cell migration, proliferation, and survival. Additionally, downregulation of GJB1 is often linked to increased melanoma cell invasiveness, reduced cell adhesion, and disruption of normal cellular signaling pathways that restrict tumor growth (75).

Our data show that GATA2, TRIM33, and PMEPA1 exhibit the most significant increase in regulatory activity, while MYBL2, NEDD4L, and STUB1 exhibit the most substantial decrease in regulatory activity. All six genes have been extensively studied in cutaneous melanoma but not in UM. Given the similarities in melanocytic tumor biology between cutaneous melanoma and UM, it is plausible that these genes play a similar role in UM. Particularly, GATA2 deficiency often due to epigenetic silencing is associated with an increased risk of melanoma (76). TRIM33 plays dual role – tumor suppressor and oncogene - in cancers (77, 78), but its relevance to melanoma remains unclear. PMEPA1 can promote tumorigenic traits in melanoma, with its expression fine-tuned by RNA-binding proteins or miRNAs to drive the tumor progression (79). MYBL2 is widely expressed in proliferating cells, and its aberrant expression contributes to tumor malignancy and poor prognosis in melanoma patients (80). MYBL2 promotes the formation of stem-like cell populations in melanoma (80). The ubiquitination activity of NEDD4 is closely linked to melanoma progression (81–83). For instance, disrupting NEDD4 activity stabilizes the tumor suppressor PTEN and inhibits melanoma cell proliferation (84, 85). Moreover, NEDD4 positively regulates expression of SOX10 (86). STUB1 regulates immune response and resistance to immunotherapy in melanoma (87, 88).

Besides protein-coding genes, we identified 17 miRNAs with significant differential activities. miR-146a-5p, miR-25-3p, and miR-34a-5p show the highest upregulated activity levels, and miR-573, miR-4426, and miR-4485-5p show the greatest downregulated activity levels. Notably, the regulatory activities of miRNAs successfully distinguished SOX10-KD cells from controls, which was not achievable using expression profiles alone, indicating that SOX10-KD reshapes miRNA regulatory networks in UM (Figure 2D).

Taken together, integrating gene expression profiles with the UM network enabled us to characterize genes’ regulatory activities in SOX10-KD UM cells, thereby identifying driver miRNAs that lack significant changes in their expression levels.

### Quantification of driver miRNAs’ regulatory activities

To better understand the role of the identified 17 miRNAs in UM, we modeled the expression patterns of these miRNAs and their target genes using NetBID2 to quantify changes in miRNAs’ regulatory activities (see Methods for details). Since miRNAs repress their target genes’ expression (49), lower expression levels of miRNA target genes indicate increased miRNA regulatory activity, while higher expression levels indicate decreased miRNA regulatory activity (Figure 3A).

**Figure 3:**
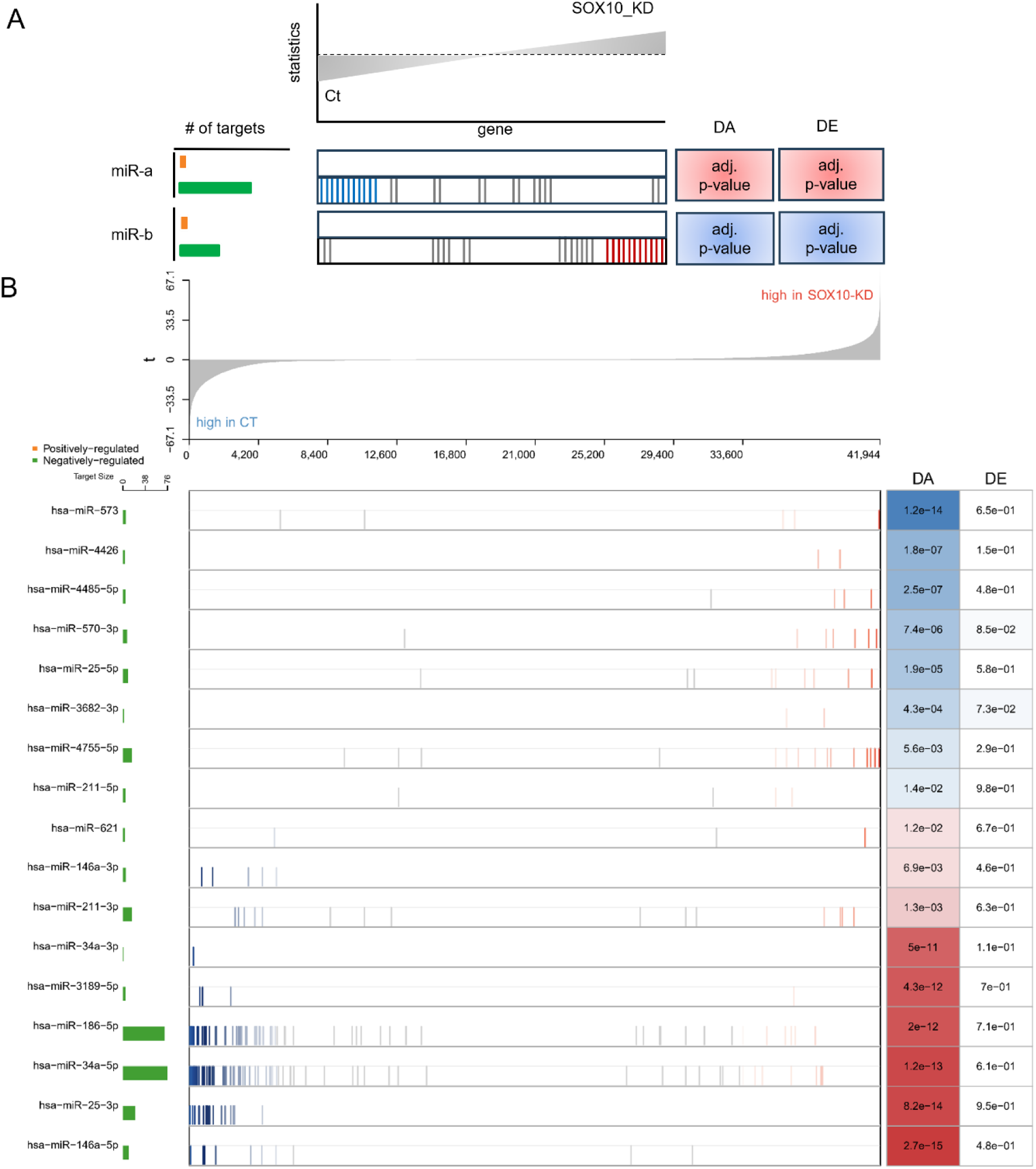
Regulatory profiles of the identified driver miRNAs. (**A**) The illustration of DA analysis results using NetBID2. The upper plot shows the distribution of target genes’ t statistics, with genes expressing higher levels in the control group positioned on the left and genes expressing higher levels in the SOX10-KD group positioned on the right. The bottom panel provides a regulatory landscape of miRNAs. Each row represents a driver miRNA and its targets’ expression profiles. Each vertical bar represents a target gene and its position that is determined by its t statistics in the upper plot. The bar in blue or red indicates whether the corresponding target gene is significantly (|z-score of t-statistics| ≥ 1.64 corresponding to p-value <= 0.10) down- or up-regulated in SOX10-KD samples, otherwise it is grey. The bar plot on the left shows the number of target genes that are positively (green) and negatively (orange) regulated by a miRNA, respectively. The orange bar is not shown (i.e. its value is zero), because miRNA-target interactions are negative regulation due to miRNA’s repressive effects on target genes’ expression levels. The boxes on the right show the DA and DE results of miRNAs. Blue and red boxes represent significantly (|z-score of t-statistics| ≥ 1.64 corresponding to p-value <=0.10; the darker color the higher z-score value) down- and up-regulated activity of miRNAs, respectively. The numbers in the boxes are the corresponding adjusted p-values (the false discovery rate approach). miR-a and miR-b are two examples whose regulatory activities have significantly increased and decreased due to the altered expression profiles of their target genes in SOX10-KD UM cells. (**B**) The results for the identified 17 driver miRNAs. The figure has the same layout as shown in the illustration. The detailed information on the driver miRNAs’ target genes can be found in SM Excel S5.

Among the identified driver miRNAs, miR-146a-5p, miR-25-3p, miR-186-5p, and miR-34a-5p have numerous target genes (from 10 to 77) in the UM network, suggesting their broad regulatory function in UM. For instance, 54 of the 77 target genes of miR-34a-5p have significantly lower expression levels in SOX10-KD UM, resulting in a strong increase in its regulatory activity (Figure 3B). The significant change in regulatory activity and the large number of target genes of miR-34a-5p suggest that it plays a pivotal role in controlling the cell cycle, proliferation, and apoptosis in SOX10-KD UM. For instance, loss of SOX10 in melanoma is known to increase p53 pathway activity (89). Our data corroborates this, showing that SOX10-KD significantly upregulates TP53 expression and activity. Concurrently, we observed a significant upregulation in both the expression and activity of miR-34a-5p. This is consistent with a known positive feedback loop where TP53 activates miR-34a, and miR-34a, in turn, reinforces TP53 activity by suppressing its negative regulator, SIRT1 (90). Interestingly, accumulation of dysfunctional or mutant TP53 protein in UM is generally associated with more invasive behavior and worse prognosis (91). Therefore, our results suggest a molecular mechanism through which SOX10 ablation in UM drives the switch from a proliferative to an invasive cellular phenotype by activating the TP53/miR-34a/SIRT1 positive feedback loop. In addition, many of miR-34a-5p’s target genes are linked to UM progression. For instance, MYC gene amplification is found in nearly 70% of UM cases and is associated with poor prognosis (92). The overexpression of BCL2 and related proteins contributes to multidrug resistance of UM cells and downregulating BCL2’s expression can partially reverse the tumors’ drug-resistant phenotype (93). CDK4/6 has been extensively studied UM, with findings showing that CDK4/6 inhibition combined with other targeted therapies such as MEK inhibitors could be a promising treatment strategy for UM (94). NOTCH1 signaling plays an important role in promoting proliferation, invasion, and metastasis in UM (95–98). Several studies show that ARF6 can regulate UM development and growth (99, 100).

On the other hand, the other miRNAs have no more than five target genes in the network, suggesting their specific regulatory function in UM (Figure 3B). For instance, two of the four target genes of miR-211-5p have significantly higher expression levels in SOX10-KD UM, leading to a decrease in its regulatory activity. The two target genes are HLA-DRB1 and IL11. HLA-DRB1 is essential for presenting extracellular peptides to CD4+ T-helper cells, which is crucial for initiating and sustaining adaptive immune responses against tumors (101). IL11 is a cytokine that primarily acts via the JAK/STAT3 pathway, which is a central driver of cell survival, proliferation, and resistance to apoptosis in cancer (102).

In summary, we identified driver miRNAs in UM and characterized their regulatory activities. Because of the complex regulatory interactions of miRNAs in the network, we need to elucidate specific miRNA-gene interactions at the pathway level to better understand their function in UM.

### Understanding driver miRNAs’ role at the pathway level

To further elaborate the role of the identified driver miRNAs in UM, we combined pathway enrichment analysis, the UM network (Figure S4), and literature search to elucidate the molecular mechanisms through which these miRNAs contribute to the effects of SOX10-KD in UM. This integrated approach enabled us to map miRNA-regulated genes to relevant biological pathways, contextualize their interactions within the reconstructed UM regulatory network, and validate their reported functions in UM biology.

The data shows that the identified miRNAs can regulate genes that are involved in cancer hallmark terms such as G2-M checkpoint and mitotic spindle of cell cycle, mTORC1 signaling, targets of E2F and MYC, and fatty acid metabolism (Figure 4). Among these miRNAs, miR-34a, miR-25, and miR-186 can regulate multiple genes covering most or even all hallmark terms. However other miRNAs, such as miR-3189 and miR-211, can regulate a few genes in specific hallmarks. In the following, we discuss the potential role of some miRNAs in UM and the complete miRNA-gene interactions in different hallmark pathways and their association with UM are summarized in SM Excel S6.

**Figure 4:**
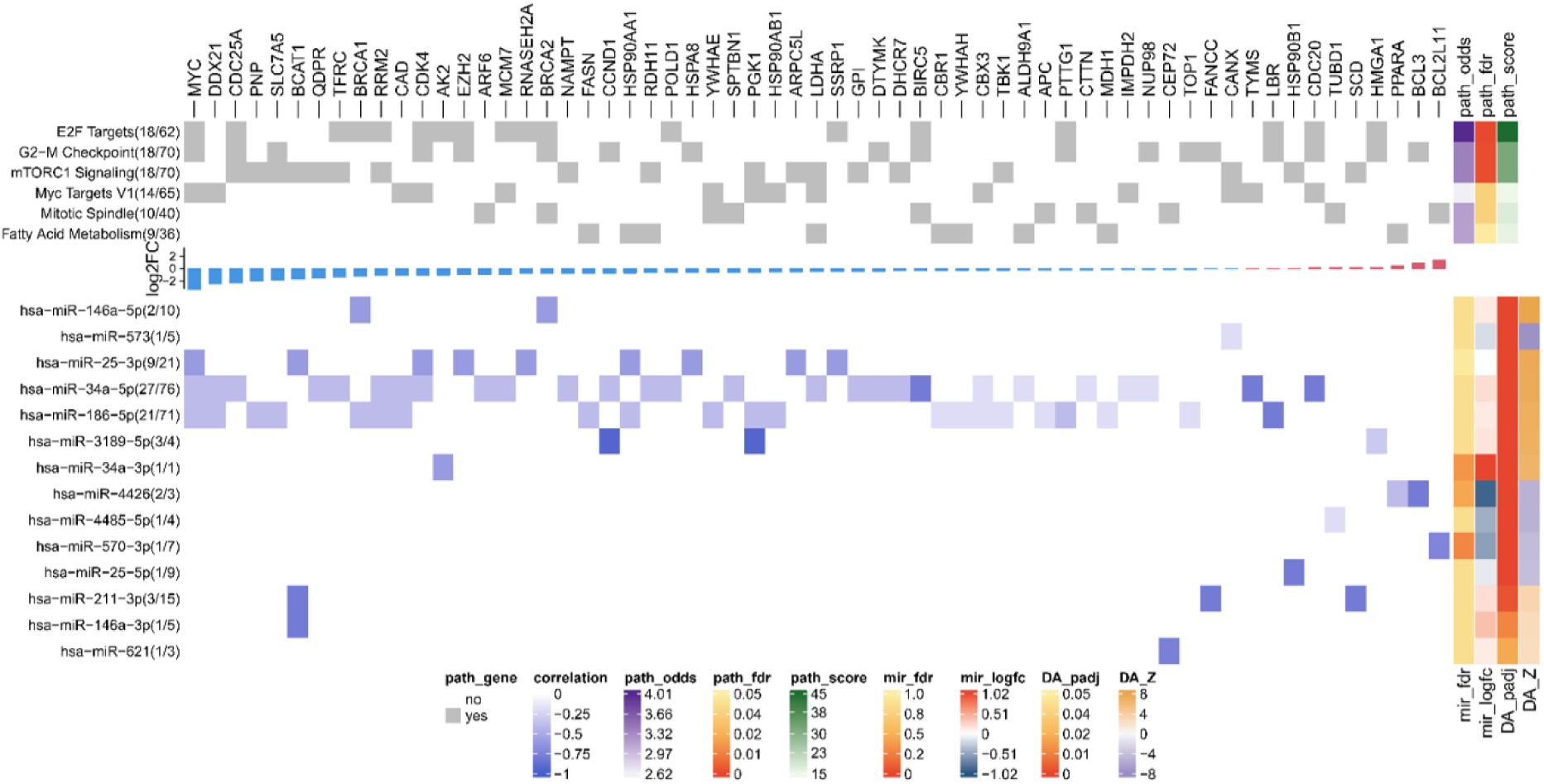
Pathway enrichment analysis results. The top heatmap shows the significantly enriched hallmark pathways and their associated genes that are targeted by the driver miRNAs. The middle bar plot shows the distribution of log2 fold-change of the genes (blue: downregulation, red: upregulation). The bottom heatmap shows the Spearman correlation coefficients between miRNAs and their target genes that are enriched in the hallmark pathways, with darker colors representing higher correlation values. The annotation bars represent the results of gene set enrichment analysis and differential gene expression and regulatory activity analysis. This includes pathways’ odds ratio, adjusted p-value, and combined score and miRNAs’ log2 fold-change and regulatory activity in Z-score normalization with the corresponding adjusted p-values.

For cell cycle, the G2-M checkpoint is frequently compromised in UM, resulting in heightened genomic instability and aggressive tumor phenotypes (103). This checkpoint disruption stems from mutations and dysregulation of critical cell cycle regulators, including GNA11, GNAQ, CDK4, and CCND1. These molecular defects facilitate hallmark cancer characteristics in UM, particularly metastatic progression and unfavorable clinical outcomes (104). Our analysis reveals that CDK4 and CCND1 are commonly targeted by miR-34a in UM, while miR-25 and miR-3189 respectively regulate these key cell cycle checkpoints (Figure 5). The enhanced regulatory activity of these miRNAs following SOX10-KD suggests that they play a pivotal role in controlling DNA damage responses and proliferative capacity in UM. On the other hand, the mitotic spindle is essential for accurate chromosome segregation during cell division. In UM, spindle dysfunction such as centrosome amplification promotes chromosomal instability, contributing to tumor progression and metastasis (105). BIRC5 encodes survivin, a chromosomal passenger complex protein essential for mitotic spindle assembly, chromosome segregation, and cytokinesis, whose overexpression in UM correlates with apoptosis resistance (106, 107) and poor prognosis (108). Our analysis shows that downregulated BIRC5 is targeted by miR-34a-5p, whose expression and activity increase following SOX10-KD, suggesting an alternative TP53-independent tumor suppressor mechanism for miR-34a in UM. BCL2L11 encodes BIM, a pro-apoptotic BCL-2 family member crucial for initiating apoptosis in response to mitotic stress or spindle checkpoint failure, and its upregulation in UM could enable elimination of genetically unstable tumor cells with spindle defects (109). We showed that upregulated BCL2L11 is targeted by miR-570-3p, which is downregulated and exhibits reduced regulatory activity following SOX10-KD, indicating the oncogenic role of this miRNA in UM progression.

**Figure 5.**
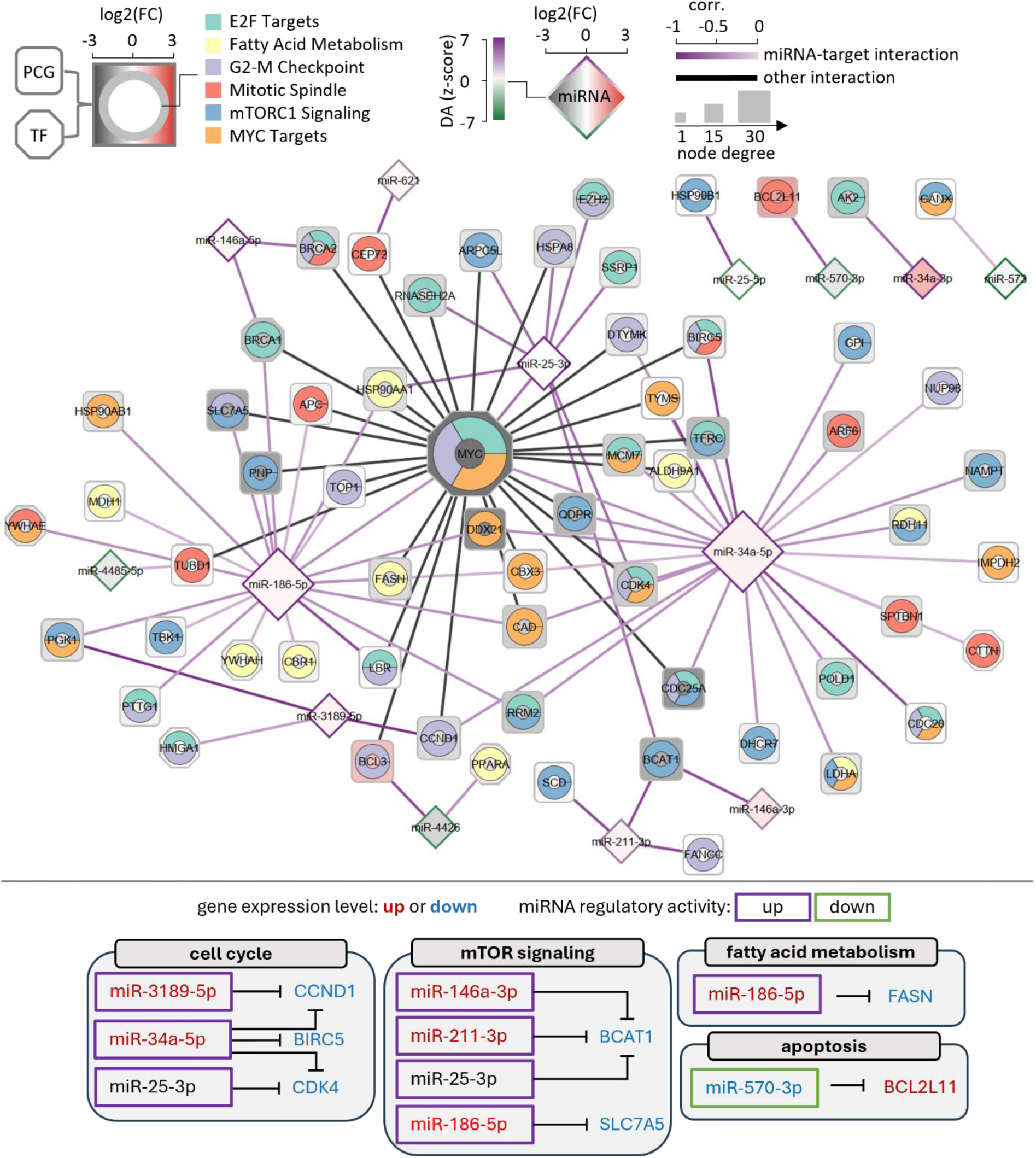
The identified key miRNA-gene interactions in the UM network. The shape of the nodes represents different molecular species, including protein-coding genes (PCGs; squares), TFs (octagons), and miRNAs (diamonds). The ring chart embedded in the PCG and TF nodes represents their associations with enriched hallmark terms. Node color represents the log2 fold-change of gene expression profiles. The edge color of the miRNAs represents changes in regulatory activities normalized using the z-score method. Node size increases with node degree. Line color represents different types of interactions, with miRNA-target interactions shown in a purple gradient quantified by correlation coefficients. Further details, including references, cell phenotypes and cancer types, of the identified microRNA-gene interactions can be found in SM Excel S6. The bottom figure summarizes the identified functional miRNA-gene interactions that are discussed in the main text. The font color indicates whether a gene is significantly up (red) or downregulated (blue). The window indicates whether a miRNA has a significant upregulation (purple) or downregulation (green) in regulatory activity.

mTORC1 signaling represents a fundamental driver of UM pathogenesis and disease progression, with its inhibition effectively reducing tumor cell proliferation, migration, and invasive potential (110, 111). SLC7A5-mediated amino acid transport sustains the metabolic requirements of rapidly dividing cells by maintaining mTORC1 activation and downstream anabolic pathways (112). Our findings show that SLC7A5 is regulated by miR-186-5p, whose activity increases following SOX10 ablation, indicating miR-186’s potential function in modulating UM tumorigenicity and metabolic reprogramming through mTORC1 activation (Figure 5). Additionally, BCAT1 catalyzes the conversion of branched-chain amino acids into α-keto acids, generating intermediate metabolites such as leucine and α-ketoglutarate that serve as mTORC1 activators (113). In cutaneous melanoma, BCAT1 depletion suppressed tumor proliferation and migration while diminishing oxidative phosphorylation (114). Our data indicates that SOX10-KD enhances the regulatory activities of miR-25, miR-211, and miR-146a, which collectively target BCAT1, suggesting their tumor suppressive functions in UM (Figure 5).

Fatty acid metabolism is crucial in UM development and progression (115), with UM cells exhibiting higher polyunsaturated fatty acid proportions than normal uveal melanocytes, contributing to abnormal proliferation and altered antioxidant status (116). FASN, which mediates de novo fatty acid synthesis and is upregulated in metastatic UM to support tumor growth (115), is targeted by miR-186-5p whose expression and regulatory activity increase following SOX10-KD, suggesting an anti-cancer role for this miRNA in regulating UM fatty acid metabolism (Figure 5).

Taken together, our analysis reveals that SOX10-KD in UM modulates multiple miRNAs that regulate hallmark pathways, including cell cycle, mTORC1 signaling, and fatty acid metabolism. These miRNAs, particularly miR-34a, miR-25, miR-186, and miR-211, show tumor suppressive potential by targeting genes involved in UM progression and metastasis. The coordinate regulation of these pathways suggests a complex regulatory network where SOX10 loss triggers compensatory tumor suppressive mechanisms through miRNA-mediated pathway modulation.

## Discussion and conclusion

This study represents the first systematic investigation into how SOX10 expression modulates miRNA activity in UM. We employed a comprehensive systems biology approach to elucidate the regulatory functions of miRNAs following SOX10-KD, integrating miRNA and mRNA expression profiles with network modeling to conduct systematic analysis of miRNA-gene interactions specific to SOX10 depletion in UM. Our network-based methodology introduces an advanced perspective by demonstrating that miRNAs can exert significant regulatory effects even without substantial fold-changes in their expression levels. This approach enabled identification of miRNAs exhibiting significant activity changes following SOX10-KD, revealing their target genes within the UM regulatory network. Functional enrichment analysis demonstrated that these target genes are significantly overrepresented in critical cancer hallmark pathways, which collectively regulate tumor progression through modulation of cell cycle and metabolism. These findings provide mechanistic insights into pathway-level miRNA functions that are influenced by SOX10’s activity in UM. While the literature supports the biological plausibility of our predicted miRNA-gene interactions and their associated molecular mechanisms, though not all specifically in UM but in other cancers, we propose experimental validation to test relevant hypotheses and validate our predictions in UM.

Many genes function as hidden drivers in cancer progression, exerting regulatory influence without exhibiting detectable genetic alterations, epigenetic modifications, or significant differential expression at the mRNA or protein levels. These hidden drivers may contribute to tumor progression through mechanisms such as post-translational modifications (117), epistatic gene interactions (118), or epigenetic tuning (119) that remain undetected by conventional analytical approaches. Traditional genome-wide differential gene expression analyses are limited in their ability to identify hidden regulators due to their reliance on the magnitude of molecular changes, such as gene expression levels. To overcome this challenge, we employed NetBID2, an algorithm that identifies prominent miRNAs using an informatics approach to infer regulatory activity changes by integrating miRNA expression levels with those of their target genes. This approach represents a paradigm shift from conventional methodologies that typically identify significantly differentially expressed genes first and subsequently construct networks from these candidates. Instead, we used a reverse strategy by initiating our analysis with a curated network that captures consensus molecular interactions for UM. This network provides a biologically informed foundation that enables more accurate and contextually relevant calculation of miRNA regulatory activity in UM, potentially revealing hidden regulatory mechanisms that would otherwise remain undetected through traditional expression-based approaches.

While our approach provides several methodological advantages, certain limitations must be acknowledged. Though effective, the method used to detect miRNA-gene interactions within the UM network may generate false positive predictions due to bioinformatics challenges, such as the minimal biological impact and non-functional conservation of miRNAs (120), as well as the intricacy of miRNA-gene interactions (121). This could potentially lead to the identification of spurious regulatory relationships. For example, miR-34a’s repression efficacy is determined by the structural and biophysical characteristics of its binding sites in the 3’ UTR of the target mRNA and AGO2, which can increase or decrease the binding strength of miRNAs (122). Seedless binding in miR-155 and miR-124 can also lead to gene repression (123). Additionally, the accuracy and reliability of NetBID2 may be constrained by our relatively small sample size, which could impact both the comprehensiveness of detected interactions and the statistical robustness of our findings. For example, the computed changes in miRNA regulatory activity in our dataset are modest in magnitude. Despite these constraints, our work presents a systematic approach for elucidating complex miRNA regulatory networks in UM. To encourage reuse and reproduction of our results, we make the generated data and code accessible in accordance with the FAIR principle (124).

## Data availability

The code and data used in this work are archived in Zenodo (https://doi.org/10.5281/zenodo.17804306).

## Declaration of AI usage

We utilized Claude (Sonnet 4.5) to enhance the clarity and readability of the paper.

## Acknowledgements

JV and XL were supported by the German Ministry of Education and Research (BMBF) through the project e:Med MelAutim [01ZX1905A]. J.V. was also supported by and the Masterplan Bayern Digital II [MED-1810-0023], the Manfred-Roth-Stiftung, Forschungsstiftung Medizin Uniklinikum Erlangen, and EU through the Horizon 2020 project CANCERNA. M.V.H. was supported by Hiege-Stiftung – die Deutsche Hautkrebsstiftung, Matthias Lackas-Stiftung, Dr. Helmut Legerlotz-Stiftung, K.L. Weigand’sche Stiftung, Else-Kröner-Fresenius Excellence Fellowship, Clinician Scientist Fellowship of the Arbeitsgemeinschaft Dermatologische Forschung (ADF). The clinical trial NCT04335890 which generated part of the sequencing data was funded by the Hasumi International Research Foundation. The funders had no role in the design of the study; in the collection, analyses, or interpretation of data; in the writing of the manuscript, or in the decision to publish the results.

## Authors’ contributions

**Chunyan Luan**: data curation; formal analysis; software; visualization; writing – original draft; writing – review and editing. **Anja Wessely**: data curation. **Zhesi Zhang:** software, visualization. **Liang Zhang**: software. **Adrian Weich:** data curation; writing – review and editing. **Christopher Lischer:** data curation. **Carola Berking**: supervision; resources; writing – review and editing. **Markus V. Heppt**: funding acquisition; writing – review and editing. **Julio Vera**: supervision; formal analysis; funding acquisition. writing – review and editing. **Xin Lai**: conceptualization; methodology; software; data curation; investigation; validation; formal analysis; supervision; funding acquisition; visualization; project administration; resources; writing – original draft; writing – review and editing.

